# Microvesicle-Mediated Tissue Regeneration Mitigates the Effects of Cellular Ageing

**DOI:** 10.1101/2022.04.13.488143

**Authors:** Nikolaos Panagiotou, Dagmara McGuiness, Armand M.G. Jaminon, Barend Mees, Colin Selman, Leon Schurgers, Paul G. Shiels

## Abstract

An ageing global population brings with it a significant burden of age-related morbidities. Recently, a novel intervention strategy to mitigate this burden has emerged, involving the use of Extracellular Vesicles (EV), comprising use of Microvesicles (MV) and Exosomes (Exo). These membranous vesicles are secreted by cells and mediate repair of cellular and tissue damage via paracrine mechanisms, involving interaction of their bioactive cargoes with stem cells. The actions of EV under normative and morbid conditions in the context of ageing remains largely unexplored. We now show that MV, but not Exo, from Pathfinder cells (PC), a putative stem cell regulatory cell type, enhance the repair of Human Dermal Fibroblast (HDF) and Mesenchymal Stem Cell (MSC) co-cultures following both mechanical and genotoxic stress. Critically, this effect was found to be both cellular age and stress-specific. Notably, MV treatment was unable to repair mechanical injury in older co-cultures, but still remained therapeutic following genotoxic stress. These observations were further confirmed in HDF and Vascular Smooth Muscle Cell (VSMC) co-cultures of increasing cellular age. In a model of comorbidity, comprising co-cultures of HDF and highly senescent Abdominal Aortic Aneurysm (AAA) VSMC, MV administration appeared to be senolytic following both mechanical and genotoxic stress, prior to enabling regeneration. To our knowledge, this is the first description of EV-based senolysis. It provides novel insight into understanding the biology of EV and the specific roles they play during tissue repair and ageing. These data will potentiate development of novel cell-free therapeutic interventions capable of attenuating age-associated morbidities and avoiding undesired effects. Ultimately, this might act as a possible intervention strategy to extend human healthspan.

## 1. INTRODUCTION

The burden of age-related morbidities is expected to increase as a direct consequence of an expanding aged global population **[1, 2]**. The capacity for tissue regeneration and wound healing diminishes with age **[3]**. Indeed, these processes are affected by the diseasome of ageing and are thus consequently impaired **[4, 5]**. Mitigating the effects of an associated diseasome of ageing **[6, 7]** is challenging and a range of therapeutic interventions have been advanced to address this problem, including senolytic drugs designed to remove senescent cells from ageing or damaged tissue, in order to restore more normative physiological function **[8-10]**. Elimination of senescent cells by a range of senolytic agents has already been demonstrated to increase lifespan and healthspan in mice **[10-13]**. However, it is unknown whether senolytic agents will function equivalently across the lifecourse or how they will function in a multi-morbid milieu **[14]**. Alternative senotherapies are also in development. These include cellular therapeutics to restore or replace damaged tissue **[15-22]**. However, many such cell-based therapies have logistical and technical hurdles associated with them, including increased cancer risk, appropriate cell selection and differentiation, targeted delivery, cost and scalability **[23, 24]**. Such therapies have had varied success in practice and their exact mechanism of action remains incompletely understood **[25]**.

Notably, recovery of tissue function following administration of cell-based therapeutics has not typically been as a result of incorporation of the cellular therapeutic directly into the regenerated tissue **[26]**, consistent with a model of paracrine-mediated tissue regeneration. Recently, this paracrine effect has been identified and characterised for Pathfinder Cells (PC), a putative stem cell regulatory cell type, that displays overlapping features with mesenchymal stem cells **[27, 28]**. Rat PC can stimulate recovery of tissue structure and function across a species barrier in both concordant and discordant xenotransplant models of acute tissue damage, including streptozotocin (SZT)-induced damage to the pancreas and ischaemia/reperfusion damage to the kidney **[27, 28]**, with further studies consistently demonstrating a paracrine mode of action in mice **[29]**.

Extracellular Vesicles (EV), including Microvesicles (MV) and Exosomes (Exo), have been proposed experimentally as plausible candidates for facilitating paracrine-mediated tissue regeneration through the transfer of bioactive cargo, including proteins, mRNA and miRNA **[30, 31]**, for both cellular and rodent models of organ damage **[32-34]**. Interestingly, several studies have provided evidence that links EV with cell proliferation **[35-37]**, cell migration **[35, 38-40]** and angiogenesis **[41, 42]** during wound healing. Nevertheless, an age-associated deterioration in wound healing capacity has been well documented **[43]** and therefore cellular ageing might impair the efficacy of EV-mediated wound healing. However, the capacity of EV to mediate wound repair has yet to be investigated in the context of cellular ageing. Additionally, the identity and nature of any EV involved in tissue regeneration *in vivo*, remains to be fully defined. Significantly, *in vivo*, in a Streptozotocin (STZ)-induced mouse model of diabetes, only MV and not Exo have been demonstrated to facilitate improved glycaemia **[29]**. The mechanistic basis of this remains unresolved. Consequently, understanding EV mechanisms of action during maintenance of organismal homeostasis and tissue regeneration throughout lifespan could prove beneficial for the development of novel senotherapies for improvement of the healthspan.

We have therefore investigated the therapeutic capacity of PC-derived EV to facilitate tissue regeneration, in the context of cellular ageing and disease. MV and Exo were investigated and cross-compared for their capacity to facilitate repair in Human Dermal Fibroblast (HDF) and Mesenchymal Stem Cell (MSC) co-cultures of increasing cellular age, following mechanical stress induced by wounding, and genotoxic stress induced by uraemic serum derived from chronic kidney disease (CKD) patients. In this tissue regeneration model, the effect of cellular ageing and underlying pathology was further investigated in HDF and human Vascular Smooth Muscle Cell (VSMC) co-cultures of both healthy and diseased Abdominal Aortic Aneurysm (AAA) origin.

## 2. RESULTS

### 2.1. Only PC-derived MV and not Exo are able to accelerate wound healing

We explored the regenerative properties of PC-derived EV to mediate wound healing in an HDF-MSC co-culture model, incorporating a scratch assay *in vitro*. The ageing characteristics of all cells used in subsequent studies, including Senescence Associated (SA) β-galactosidase, Cytoplasmic Chromatin Fragments (CCF), CDKN2A and CDKN1A expression, are shown in **Figure S1**. All ageing biomarkers demonstrated an increase with increasing cellular age. For the purposes of this investigation, co-cultures comprising HDF and MSC were employed to provide a basic surrogate for the tissue microenvironment **[44]**. Specifically, we investigated and cross-compared MV and Exo for their therapeutic efficacy to facilitate wound repair. MV and Exo were isolated from cell culture media of rat PC, through a series of sequential centrifugation and ultrafiltration steps **[29]**. They were then characterised by their size, surface markers and miRNA cargo. The MV isolates ranged between 0.1 and 1 μm in size, as expected **[45]**, while the Exo were undetected with flow cytometry, as a consequence of their significantly smaller size and the triggering threshold of the instrument **(Figure S2A)**. Integrin β1, CD40, Rab5b and CD63 were also found to be expressed on the MV isolates, while CD9 was expressed only in the Exo preparations **(Figure S2B)**, in agreement with previous observations **[29, 46, 47]**.

We monitored wound repair with Real Time Cell Analysis (RTCA) proliferation assays over a three-day period following wounding. Administration of Exo directly after wounding had no significant beneficial effect on wound healing, while addition of MV significantly enhanced the efficiency of cells to proliferate in response to wounding. Additionally, MV-treated co-cultures were capable of complete wound closure, whereas Exo treatment and non-treated controls failed to achieve this **(Figure 1A and 1B)**.

**Figure 1:**
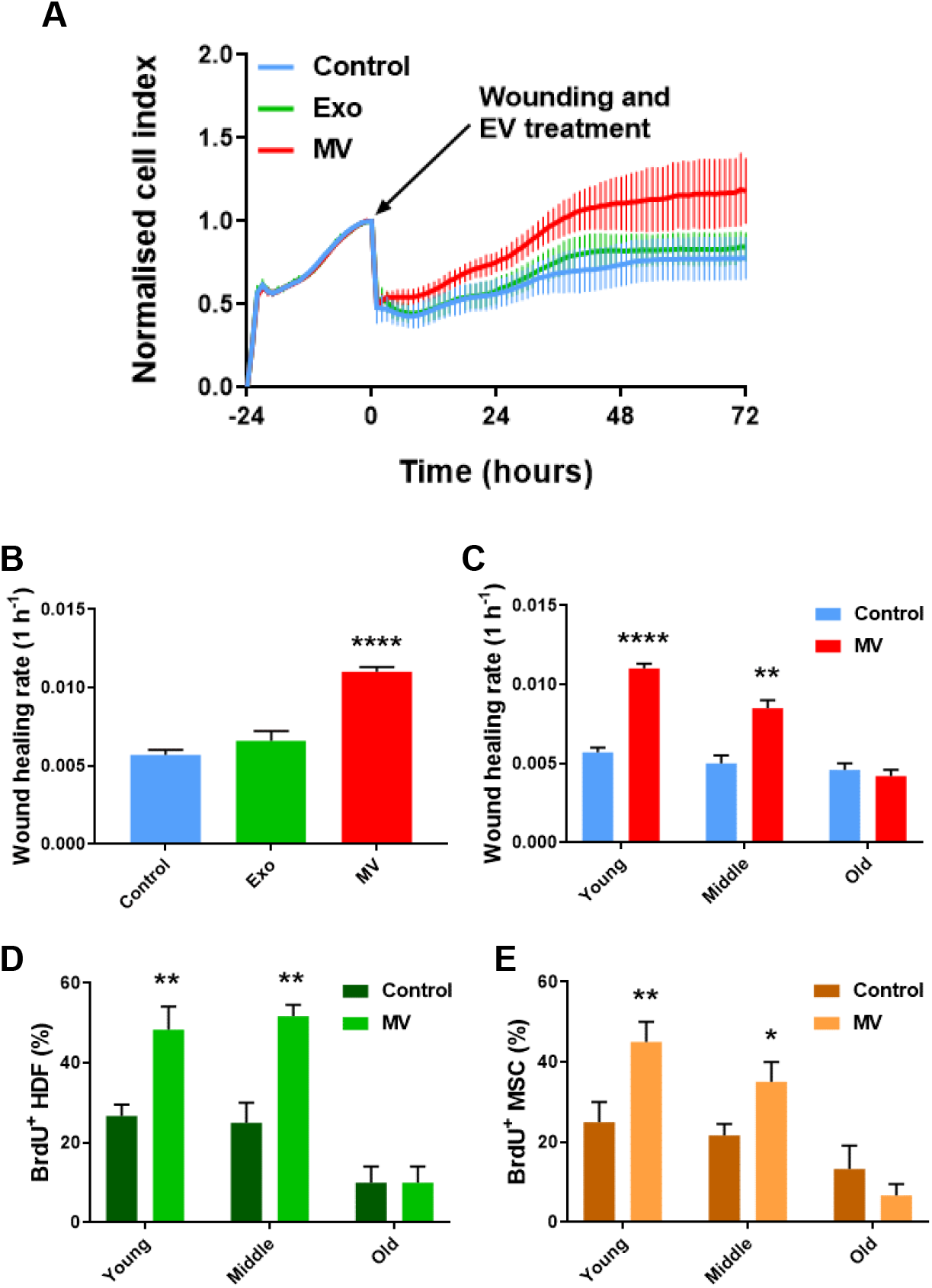
MV treatment improves cellular proliferative capacity following wounding, while old cellular age inhibits the MV beneficial effect. (A)RTCA proliferation assay of a wound healing assay on a young HDF-MSC co-culture. Data are presented as mean ± SD (n=6). (B)Wound healing rate of young HDF-MSC co-cultures treated with MV and Exo, from 0 to 72 hours. Data are presented as mean ± SD; One-way ANOVA with Tukey’s test (N=3). (C)Wound healing rate of HDF-MSC co-cultures of increasing cellular age, after wounding and under MV treatment. Data are presented as mean ± SD; t test (N=3). (D)Percentage of BrdU^+^ HDF of increasing cellular age, after wounding and under MV treatment. Data are presented as mean ± SD; t test (N=3). (E)Percentage of BrdU^+^ MSC of increasing cellular age, after wounding and under MV treatment. Data are presented as mean ± SD; t test (N=3). *p < 0.05, **p ≤ 0.01, ****p ≤ 0.0001.

### 2.2. MV-mediated wound healing capacity declines in HDF-MSC co-cultures with increased cellular age

Individual cell types within co-cultures were characterised by age (passage number), into young (passage 1-4), middle (passage 8-12), and old (passage 16-20) respectively. Their proliferative capacity was then determined by RTCA slope analysis in response to mechanical stress (scratch wounding). Both young and middle age co-cultures displayed increased growth slopes following addition of MV and were able to completely close the wound over the time course of the experiment. However, older co-cultures were refractory to MV treatment and were not able to fully close the wound over the 72-hour time period of the experiment, equivalent to the untreated controls **(Figure 1C)**. Further analysis indicated that unassisted wound closure rate decreased with increased cellular age **(Figure S3A)** and this cellular age effect resulted in a significantly more rapid decrease in MV-mediated wound healing **(Figure S3B and S3C)**. 5-bromo-2’-deoxyuridine (BrdU) staining was then employed to further validate that any MV associated increase in wound closure rate was due to enhanced cell proliferation. The staining was performed individually for HDF and MSC mono-cultures. MV administration resulted in increased number of actively proliferating HDF and MSC in young and middle age cultures. No difference was observed between treated old HDF, treated old MSC, and controls **(Figure 1D and 1E)**.

To further understand the potential processes through which MV facilitate wound healing in HDF-MSC co-cultures, time-lapse fluorescence microscopy was utilised to observe migration of these cells into the wound site, in response to MV treatment during wound closure. HDF and MSC were differentially stained with CFSE and PKH26 respectively and observed at 24-hour intervals over a 3-day period. In the presence of MV, significantly more cells migrated into the site of mechanical damage, and this response was observed from as early as 24 hours post wounding. The positive effect of MV treatment also resulted in a higher number of cells at the 48 and 72 hours **(Figure 2A and 2B)**. Increasing cellular age diminished the migratory efficacy of cells during MV-mediated wound healing. More precisely, in middle cellular age co-cultures, a 24-hour delay was observed in MV-mediated cell migration, comparing to young co-cultures **(Figure 2C)**. Old age co-cultures were non-responsive to the treatment **(Figure 2D)**.

**Figure 2:**
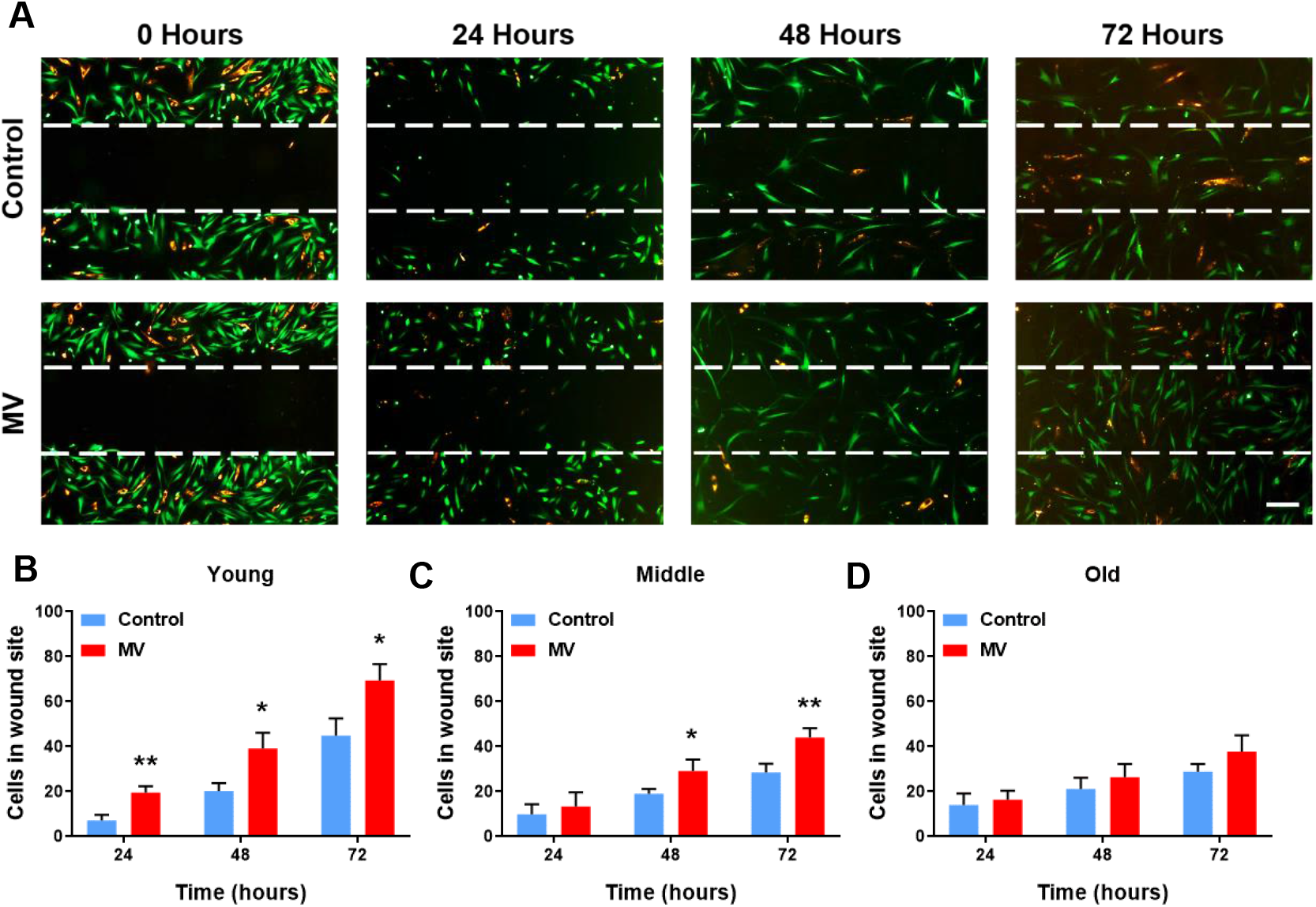
MV improve the migration of HDF and MSC in the wound site but in an age-dependent manner. **(A)** Time-lapse fluorescence microscopy of a wound healing assay in a young HDF-MSC co-culture, treated with MV. Wounding and MV administration took place at time point 0. HDF labelled with CFSE (green) and MSC with PKH26 (orange). Magnification 50x, scale bar 100 μm. Effect of MV administration on cell migration into the wound site; 24, 48 and 72 hours after wounding in **(B)** young, **(C)** middle and **(D)** old HDF-MSC co-cultures. Data are presented as mean ± SD; t test (N=3). *p < 0.05, **p ≤ 0.01.

The response of each of the two co-cultured cell types to MV treatment during wound closure was then studied individually. MV addition had a positive effect on young HDF, increasing cell migration into the wound site from 24 hours onwards **(Figure S4A)**. This response was delayed in middle cellular age HDF and significantly more HDF migrated into the wound site from 48 hours onwards **(Figure S4B)**. Finally, no differences were detected in the number of HDF that migrated in the site of damage at old cellular age **(Figure S4C)**. Similarly, young MSC responded in a positive manner to MV administration and migrated in the wound site from at 24, 48 and 72 hours **(Figure S4D)**. In middle cellular aged MSC, as in HDF, the MV positive effects, in terms of cell migration, were only apparent from 48 hours onwards **(Figure S4E)**. Finally, there were no observable differences in the number of cells that migrated into the site of damage, between old MV-treated MSC and controls **(Figure S4F)**.

Overall, these data suggest that MV have the capacity to enhance the migration rate of both HDF and MSC in response to mechanical stress and repair wounds *in vitro*. This effect occurred in a time-dependent manner that was negatively affected by cellular ageing. Explicitly, the therapeutic effect was delayed in middle-age and not observed in old co-cultures. Hence, these observations further validate that cellular ageing has a negative impact on MV-mediated repair of mechanical stress. Interestingly, MV did not increase the proliferative or migratory capacity in control unwounded HDF-MSC co-cultures of increasing cellular age **(Figure S5)**, indicating that they enhance cell proliferation and migration only in the presence of cellular damage.

### 2.3. Cellular age does not impact MV efficacy under morbid conditions

As premature ageing is a common feature of the diseasome of ageing, including CKD **[48, 49]**, we sought to determine if features of morbidity (i.e., uraemia) affected MV efficacy in the context of cellular age. Co-cultures comprising HDF and MSC were employed to assess the regenerative properties of PC-derived MV and Exo in facilitating repair of genotoxic stress caused by uraemia derived from CKD stage 3 patients. During the first 24 hours following the addition of uraemic serum, there was a distinct period of decreased cell growth. Interestingly, the RTCA data suggested that MV administration resulted in a significant decrease in cell growth, in comparison to both control and Exo treatment, during this period of genotoxicity **(Figure 3A and S6A)**. Nevertheless, over the subsequent 48 hours, an increase was observed in cell growth for the MV-treated samples. This time window was termed as *healing period* because the cells were able to recover from the genotoxic stress and proliferate. During the healing period, the data indicated that MV addition had a positive effect on cell growth, while the control cell index did not significantly increase. In the control, this is indicative of a cytostatic effect in response to the presence of uraemic serum. Conversely, MV were subsequently able to facilitate a positive response and cell growth significantly surpassed that of the control by the end of the experiment **(Figure 3A)**. During the healing period, slope analysis indicated that administration of MV significantly enhanced the rate of cell growth **(Figure 3B)**. This was also true for co-cultures of all cellular ages that were investigated. Hence, the MV-associated therapeutic effect was not only recorded in young and middle cellular age mixed cultures, as it was the case with wound healing assays, but also in co-cultures of old cellular age **(Figure 3C)**. Furthermore, the increased MV-mediated cell loss during the genotoxicity period was also observed at all cellular ages **(Figure S6B)**. Administration of Exo directly after induction of genotoxicity had a protective effect during the initial 24-hour genotoxicity period. The Exo treated samples retained a higher cell index and significantly reduced cell loss compared to both untreated controls and the MV-treated samples **(Figure 3A and S6A)**. Nevertheless, there was no significant Exo-associated effect during the healing period. The cell growth rate remained at similar levels to that of the untreated controls and at significantly lower levels to MV-treated samples **(Figure 3B)**.

**Figure 3:**
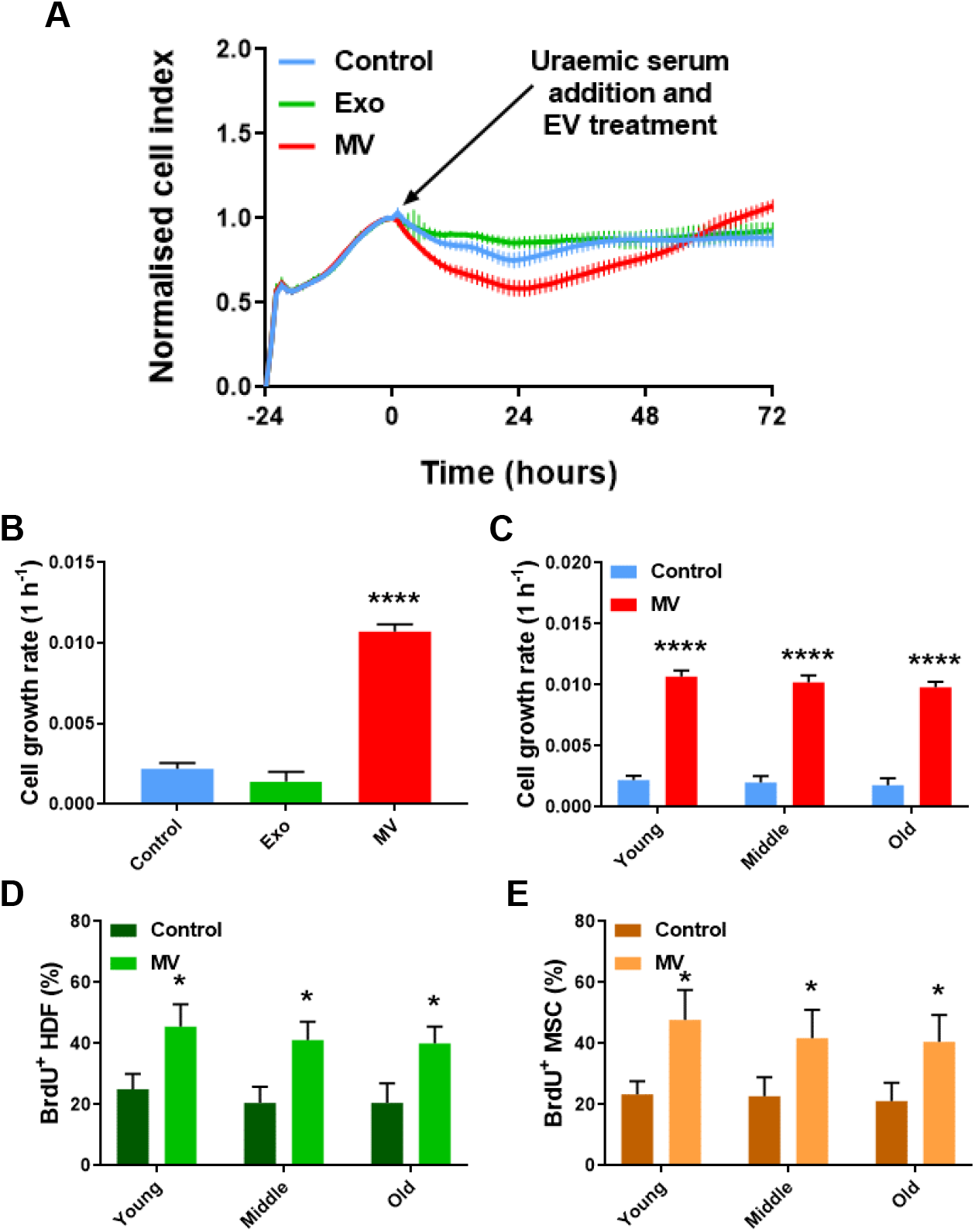
MV treatment improves cellular proliferation following uraemic stress irrespective of cellular age. (A)RTCA proliferation assay of a genotoxicity assay on a young HDF-MSC co-culture. Data are presented as mean ± SD (n=6). (B)Cell growth rate of young HDF-MSC co-cultures treated with MV and Exo, from 0 to 72 hours. Data are presented as mean ± SD; One-way ANOVA with Tukey’s test (N=3). (C)Cell growth rate of HDF-MSC co-cultures of increasing cellular age, after genotoxic stress and under MV treatment. Data are presented as mean ± SD; t test (N=3). (D)Percentage of BrdU^+^ HDF of increasing cellular age, after genotoxic stress and under MV treatment. Data are presented as mean ± SD; t test (N=3). (E)Percentage of BrdU^+^ MSC of increasing cellular age, after genotoxic stress and under MV treatment. Data are presented as mean ± SD; t test (N=3). *p < 0.05, **p ≤ 0.01, ****p ≤ 0.0001.

These data are consistent with MV treatment being a senotherapeutic. MV may thus act as a putative senolytic agent in these assays, initially enabling removal of senescent cells resulting from morbid conditions and subsequently promoting repair of the damaged tissue. This hypothesis builds upon previous studies that have demonstrated the ability of apoptotic cells to induce proliferation in neighbouring cells through compensatory proliferation signalling **[50, 51]**, mediated by the production and release of MV that stimulate proliferation **[52]**. Other studies have also indicated that MV are responsible for inducing apoptosis and cell proliferation following stress stimuli **[53-55]**. Thus, PC-derived MV may play a key role in assisting and enhancing compensatory proliferation signalling following uraemic stress, initially inducing cell loss and subsequently increasing cell proliferation.

### 2.4. MV treatment in the context of cellular abnormality

Senotherapeutic strategies, such as those employing senolytic drugs, enable removal of senescent cells for the purposes of improving age-related physiological functional capacity, as a means of extending healthspan **[9, 11, 56]**. Senolytic activity, however, could have an adverse consequence in some situations, as a result of the removal of senescent cells whose presence is essential for maintaining tissue integrity. For instance, human Vascular Smooth Muscle Cells (VSMC) in aneurysmal tissues have been identified to comprise populations of senescent cells **[57-62]** and their removal could hypothetically result in undesirable aneurysmal rupture with dire physiological consequences. Abdominal Aortic Aneurysm (AAA) is an important health problem with a prevalence of up to 9% in adults over the age of 65 **[63-65]**, thus the risk to AAA in response to senotherapies merits further investigation. Here, we assessed the effect of MV administration to mediate repair in the presence of comorbidity, associated with increased levels of cellular senescence. For this reason, AAA VSMC were compared with healthy VSMC derived from the same patient, and co-cultured with HDF, for their ability to repair mechanical and genotoxic stress *in vitro*, in response to MV treatment and with respect to cellular ageing.

For normal (i.e., non-aneurysmal) VSMC co-cultures, addition of MV significantly enhanced the efficiency of cells to proliferate in response to wounding, in comparison to the non-treated controls in young and middle-aged co-cultures **(Figure 4A and 4B)**. However, the addition of MV did not have an observable effect on wound closure rate in old mixed cell cultures. These co-cultures were refractory to MV treatment during the 72 hours of the experiment **(Figure 4B)**. Actively proliferating healthy VSMC (BrdU positive), were quantified and it was observed that MV administration resulted in complete wound closure and increased number of actively proliferating cells, in comparison to the untreated controls. This was only true for young VSMC, while middle and old age VSMC showed no difference from controls **(Figure 4C)**.

**Figure 4:**
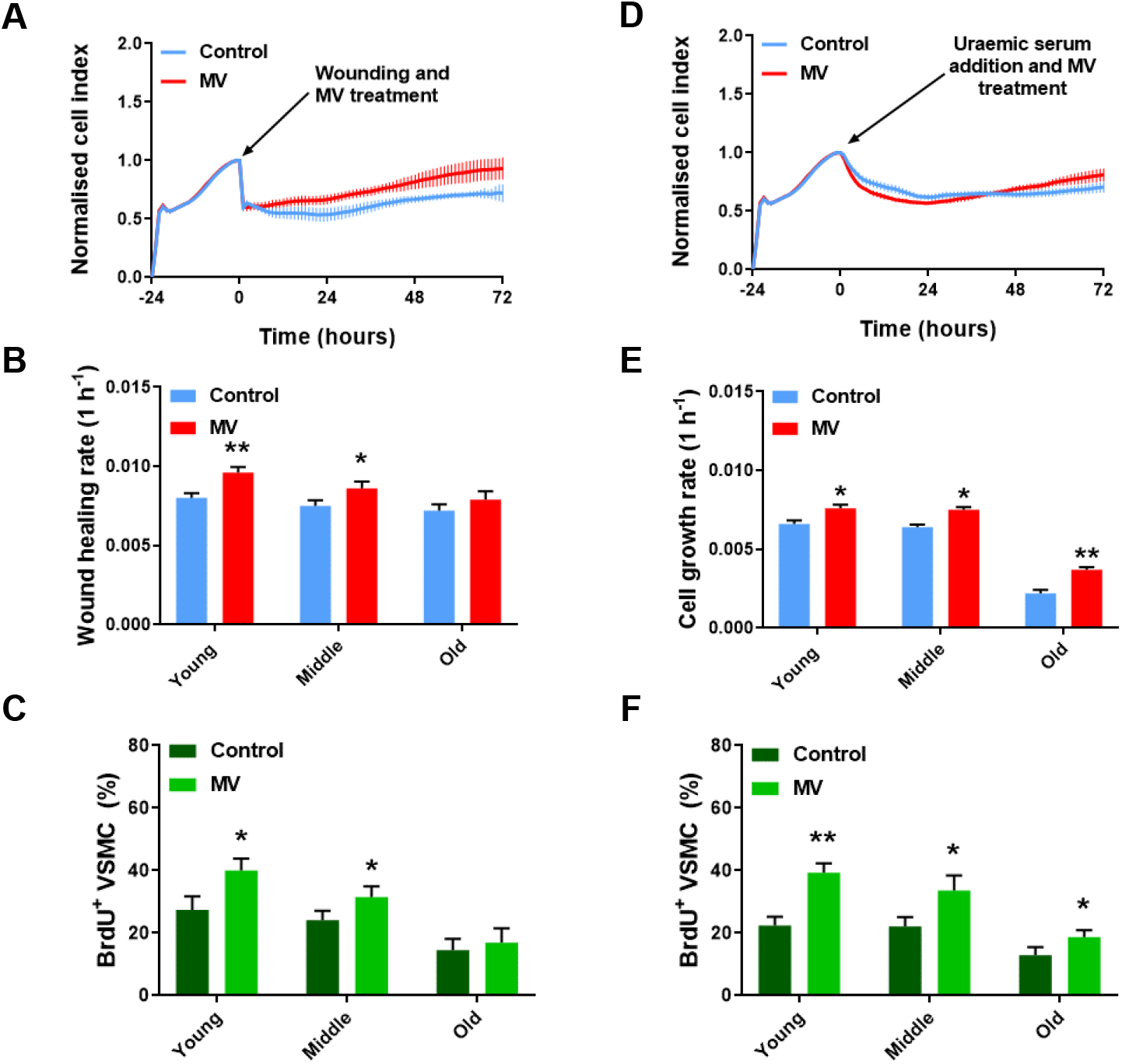
MV improve the proliferation of HDF-VSMC co-cultures in response to wounding and uraemia in a stress and age-dependent manner. (A)RTCA proliferation assay of a wound healing assay on a young HDF-VSMC co-culture. Data are presented as mean ± SD (n=6). (B)Wound healing rate of HDF-VSMC co-cultures of increasing cellular age, after wounding and under MV treatment. Data are presented as mean ± SD; t test (N=3). (C)Percentage of BrdU^+^ VSMC of increasing cellular age, after wounding and under MV treatment. Data are presented as mean ± SD; t test (N=3). (D)RTCA proliferation assay of a genotoxicity assay on a young HDF-VSMC co-culture. Data are presented as mean ± SD (n=6). (E)Cell growth rate of HDF-VSMC co-cultures of increasing cellular age, after genotoxic stress and under MV treatment. Data are presented as mean ± SD; t test (N=3). (E)Percentage of BrdU^+^ VSMC of increasing cellular age, after genotoxic stress and under MV treatment. Data are presented as mean ± SD; t test (N=3).

In uraemic genotoxicity assays, MV administration resulted in a significant decrease in cell index (thus reduced cell growth), relative to controls during the genotoxicity period from 0 to 24 hours after uraemic serum addition. Following the genotoxicity period, an increase was detected in cell index in the MV-treated samples. Specifically, the cell index surpassed that of the untreated controls. Hence, elevated cell growth during the healing period from 24 to 72 hours of the experiment was attributed to MV action, with these cells able to recover from the genotoxic stress and proliferate. The untreated controls were unable to recover from the same genotoxic stress and after an initial reduction in cell index (0-24 hours), cell growth remained unchanged from 24 hours until the end of the experiment **(Figure 4D)**. During the initial 24-hour genotoxicity period, slope analysis indicated that presence of MV significantly reduced cell index and cell growth relative to that of the controls. Consequently, MV administration following genotoxicity, appeared to increase cell loss in comparison to the non-treated controls. Cell loss was elevated as the cellular age of the co-cultured cells increased **(Figure S7A)**. This is consistent with senescent cell accumulation with cellular age and their subsequent removal following repair after genotoxic damage. During the healing period, from 24 to 72 hours, the slope analysis indicated that administration of MV significantly enhanced the rate of cell growth in all age classes, in comparison to the controls **(Figure 4E)**. However, the rate of cell growth associated with old cellular age was lower than that of young and middle cellular age co-cultures. Additionally, 72 hours post addition of uraemic serum, it was observed that MV administration enhanced the number of healthy actively proliferating BrdU positive VSMC compared to untreated controls. This observation remained unchanged, irrespective of cellular age in the assay **(Figure 4F)**.

In AAA VSMC co-cultures, addition of MV significantly reduced the efficiency of cells to proliferate in response to wounding, in comparison to the controls **(Figure 5A)**. Slope analysis from 0 to 72 hours indicated that the proliferative capacity of young cellular aged co-cultures, in response to mechanical stress, was significantly decreased after addition of MV. Additionally, these co-cultures retained a negative slope over the 72-hour period and did not display signs of recovery. These data suggest that addition of MV resulted in cell loss. However, middle cellular age co-cultures were able to respond positively to MV administration and the rate of wound repair was significantly enhanced, in comparison to the controls, while addition of MV in old co-cultures did not have an observable effect on the rateof wound closure **(Figure 5B)**. Three days post wounding, BrdU staining indicated that MV administration resulted in reduced cell proliferation in young AAA VSMC cultures. Nonetheless, complete wound closure and increased number of actively proliferating cells was observed with middle age AAA VSMC cultures, in comparison to untreated controls. We hypothesised that through cell culture and continuous passaging of cells, non-senescent cell selection hapenned naturally and thus senescent cells numbers dropped, as confirmed by the significantly lower levels of SA β-gal and CCF that were observed in middle age AAA VSMC, comparing to young highly senescenct AAA VSMC **(Figure S1A and S1B)**. Finally, MV-treated old cellular aged cultures from AAA derived cells were indistinguishable from controls **(Figure 5C)**, indicating that proliferative senescence occurred **(Figure S1)** and hence increased cellular age was a factor in cells becoming refractory to treatment.

**Figure 5:**
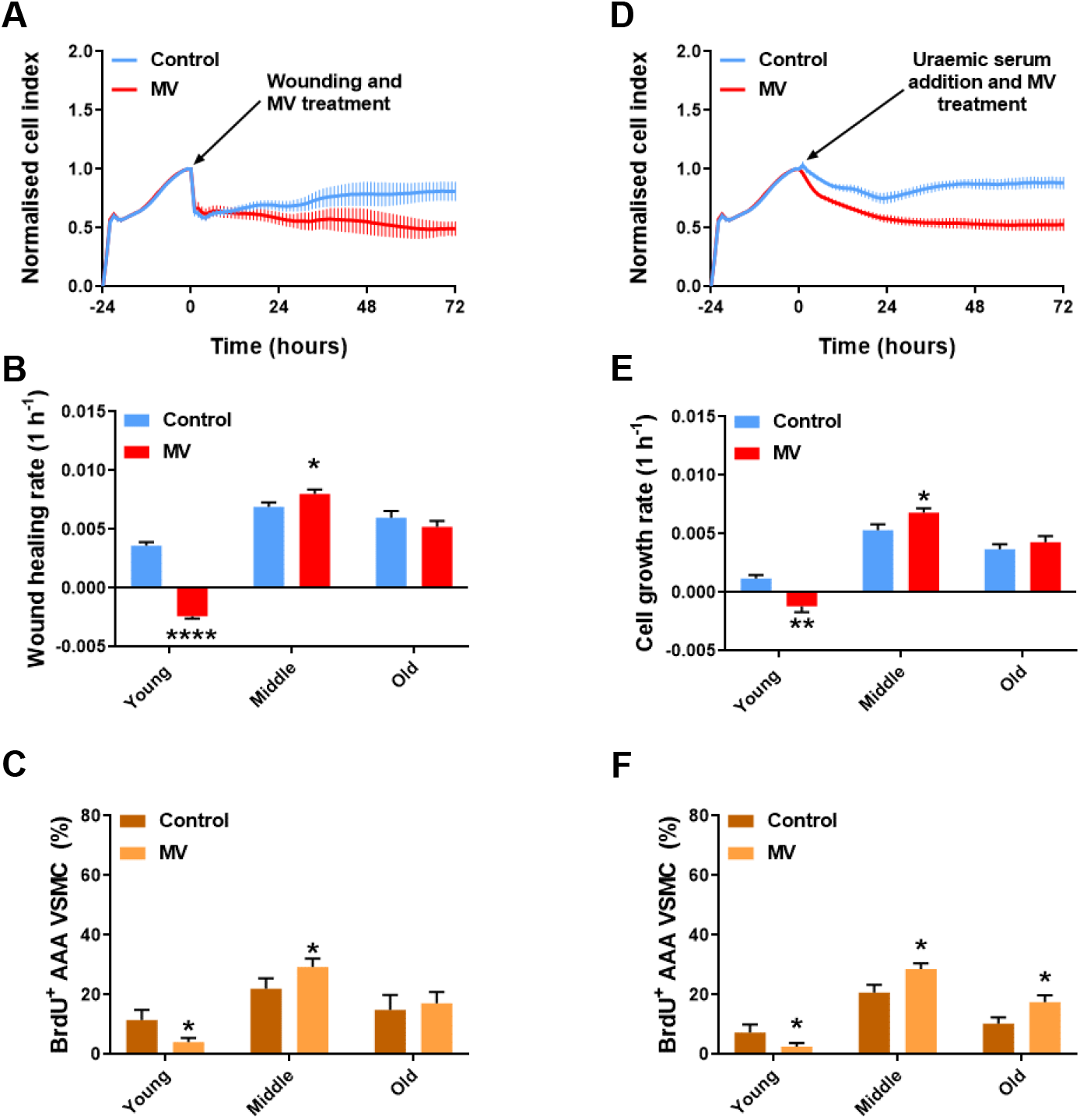
HDF-AAA VSMC co-cultures in response to MV treatment following wounding and uraemia. (A)RTCA proliferation assay of a wound healing assay on a young HDF-AAA VSMC co-culture. Data are presented as mean ± SD (n=6). (B)Wound healing rate of HDF-AAA VSMC co-cultures of increasing cellular age, after wounding and under MV treatment. Data are presented as mean ± SD; t test (N=3). (C)Percentage of BrdU^+^ AAA VSMC of increasing cellular age, after wounding and under MV treatment. Data are presented as mean ± SD; t test (N=3). (D)RTCA proliferation assay of a genotoxicity assay on a young HDF-AAA VSMC co-culture. Data are presented as mean ± SD (n=6). (E)Cell growth rate of HDF-AAA VSMC co-cultures of increasing cellular age, after genotoxic stress and under MV treatment. Data are presented as mean ± SD; t test (N=3). (F)Percentage of BrdU^+^ AAA VSMC of increasing cellular age, after genotoxic stress and under MV treatment. Data are presented as mean ± SD; t test (N=3). *p < 0.05, **p ≤ 0.01, ****p ≤ 0.0001.

In genotoxicity assays, addition of MV had a significant negative impact on cell growth **(Figure 5D)**. Slope analysis from 0 to 24 hours indicated that the proliferative capacity of young co-cultures, in response to genotoxic stress, was significantly decreased after addition of MV **(Figure S7B)**. Interestingly, young co-cultures retained a negative slope over the 72-hour period and did not display any signs of recovery, while middle aged co-cultures that were treated with MV were able to respond positively to MV administration and recover by the end of the experiment, through a significant increase in cell growth compared to the untreated controls. No effect was observed in old cellular age co-cultures **(Figure 5E)**. In AAA VSMC mono-cultures, quantification of BrdU positive cells revealed that MV decreased the ability of cells to proliferate in the presence of uraemic serum, at young cellular age. Nevertheless, MV administration was associated with an increased percentage of actively proliferating cells in middle and old cellular age AAA VSMC **(Figure 5F)**.

## 3. DISCUSSION

Due to population ageing, there is an ongoing demographic shift which will lead to a significant increase in the number of elderly individuals around the world **[66]**. This represents a major societal and global health challenge with massive economic implications, due to the elevated burden of age-related morbidities that is estimated to increase accordingly **[1, 2, 67]**. Thus, measures are sought to improve the healthspan and increase the number of years of healthy old age **[32]**. The ageing trajectory can be altered by genetic, dietary and pharmacological interventions, as demonstrated in model organisms **[68-71]**. However, the likelihood that these interventions can be directly translatable in humans is limited by unwanted side-effects or because they are impractical to undertake **[68]**. Consequently, there is need to identify novel, safe and practical interventions that ultimately improve late-life health in humans.

Therapeutic strategies that activate relevant repair processes in severely or terminally non-functional tissue are arguably the best approach for dealing with age-related pathophysiology **[32]**. Cell-based therapies, using a number of different cell types, such as induced pluripotent stem cells (iPSC) **[15, 16]**, Mesenchymal Stem Cells (MSC) **[17-20]**, very small embryonic-like stem cells **[21, 22]**, Pathfinder Cells (PC) **[27, 28, 72]** and others, are actively being developed **[73, 74]**. Nevertheless, they are beset by logistical, technical and ethical hurdles **[23, 24]**. These therapies have had varied success and their exact mechanism of action remains poorly understood. Critically, recovery of tissue function by particular cellular therapies, has typically been suggested to occur without incorporation of the cellular therapeutic directly into the tissue being investigated **[26]**. These observations are consistent with a paracrine-mediated tissue regeneration model. Extracellular Vesicles (EV) have been proposed experimentally as strong candidates for mediating such a paracrine effect, as they are involved in cell-to-cell communication **[30, 31]**. EV have previously been explored for their capacity to facilitate paracrine-mediated tissue regeneration in cellular and rodent models of organ damage **[32, 33, 45, 75]**. Such therapeutic strategies that enable severely or terminally non-functional tissue to activate relevant repair processes are arguably the best approach for dealing with age-related loss of physiological functional capability **[32]**.

In this study, we have demonstrated that only the Microvesicle (MV) component of EV derived from PC demonstrate regenerative capability and therapeutic efficacy to initiate and enhance a repair process *in vitro*, following both wounding and uraemic serum-induced genotoxicity. This observation is consistent with a previous report linking PC-derived MV with tissue repair *in vivo*, in a model of Streptozotocin (STZ)-induced diabetes **[29]**. While MV were shown to be therapeutically superior, Exosome (Exo) treatment was associated here with an anti-apoptotic effect following uraemic serum addition. Although, Exo action alone was not sufficient to enhance repair and their effects were short-term, acting together with MV might prove an effective therapeutic strategy that is worth exploring in the future.

Additionally, our *in vitro* data indicate that MV significantly enhance wound repair, associated with increased proliferation and cellular migration of both HDF and MSC into the wound site following mechanical injury, but this effect was negatively affected by cellular ageing. Thus, an interplay has been indicated to exist between MV-mediated regeneration and ageing and appears context dependent. Specifically, ageing appeared to interplay and negatively impact MV-mediated repair of mechanical, but not genotoxic, stress. This suggests that MV-facilitated repair processes deteriorate with age at different rate with respect to type of stress that induces cellular repair. Notably, MV did not increase proliferative or migratory capacity in unwounded controls, irrespective of cellular age. Hence this effect is only observed in the presence of cellular damage.

Our data are consistent with a scenario where MV can act as a senotherapeutic and senolytic agent, promoting cell loss and enabling restoration of physiological homeostasis in damaged tissue. This is in keeping with previous studies that demonstrated ability of MV derived from apoptotic cells to induce proliferation in neighbouring cells through compensatory proliferation signalling **[52-55]**. This is also consistent with PC-derived MV enabling compensatory proliferation signalling in the presence of uraemic stress. In genotoxicity assays, we observed that MV improved cellular growth repsonses and enhanced recovery of aged cells, unlike when grown under normative conditions, or when subject to mechanical stress. Notably, MV addition initially resulted in increased cell loss before enhancing cell growth, indicative of a senolytic effect where senescent cell removal is enabled.

This was further validated in AAA VSMC, which were highly senescent as others have also demonstrated for both abdominal and thoracic aortic aneurysms **[57-60, 76]**. MV administration lead to significantly increased cell loss in the presence of both wounding and uraemic serum at young cellular age. In contrast, MV administration was associated with an increased percentage of actively proliferating cells following stress in both middle and old AAA VSMC, where senescent cell presence was lower in comparison to young cellular age. These findings suggest that MV-associated putative senolytic effect could have negative implications in the presence of underlying pathology, such as AAA, that associates with elevated levels of cellular senescence and in the presence of a secondary stressor (i.e. wounding or uraemia). As others have discussed recenltly with regards to senotherapies, a cautionary approach should be adopted to avoid adverse events [77], such as the one reported in this study. Overall, highly senescent populations of cells which have resulted from comorbidity, such as AAA VSMC, are less capable of responding to subsequent stress, while MV-mediated regeneration demonstrates a deleterious effect, causing cell removal and reduced capacity for tissue repair. These observations remain to be further investigated in animal models of aneurysm and comorbid conditions, where senescent cells play an essential role in maintaining tissue homeostasis and their removal could putatively lead to undesirable consequences for the organism, such as aneurysm rupture. Additioanlly, our data inidcate that exploration of MV payloads are essential to understanding the mechanistic basis of our observations and thus merit investigation.

### 3.1. Conclusion

Under normative growth conditions, MV appear inert with respect to effects on cell proliferation and migration. However, they are therapeutically efficacious and show an age dependent effect in the presence of mechanical wounding, but not in the presence of genotoxicity. Explicitly, MV treatment repaired middle aged co-cultures at a slower rate and was unable to repair mechanical injury in old co-cultures, nevertheless remained therapeutically efficacious following genotoxic stress. Thus, MV-mediated repair interplays with ageing in a stress-specific manner. The reasons for this are unclear. There is a significant paucity of information on the role of EV in ageing and our data help address some aspects of this, specifically with respect to cell type (primary versus regenerative) and how these behave in the face of exposome stressors (physical versus physiological stress). One benefit of such a therapeutic approach is that it minimises a series of risks associated with cell-based interventions, including reduced risk of oncogenesis, decreased risk of small vessel blockage, less variability in the therapeutic entity, and easier quality assurance and quality control for getting to clinical usage, as well as scalability, logistics and cost.

## 4. METHODS

### 4.1. Cell culture

The cells used in this study were adherent and cultured in Corning® T75 flasks (Merck, UK) at 37 °C, in a humidified incubator atmosphere maintained at 5% CO2. For passaging, the cells were washed twice with phosphate buffered saline (PBS) (Thermo Fisher Scientific, UK) and trypsinised at 37°C for 5 minutes with Trypsin-EDTA (0.05%) (Thermo Fisher Scientific, UK). Complete cell media were used to deactivate the Trypsin and the cells were then seeded at a 1:3 ratio in new Corning® T75 flasks.

#### Pathfinder Cells (PC)

Rat PC of pancreatic origin were cultured in CMRL 1066 medium with no glutamine (Thermo Fisher Scientific, UK), supplemented with 10% heat inactivated fetal bovine serum (FBS) (Thermo Fisher Scientific, UK), 1% penicillin-streptomycin (Thermo Fisher Scientific, UK), 1% Amphotericin B (fungizone®) (Thermo Fisher Scientific, UK) and 1% GlutaMAX™ (Thermo Fisher Scientific, UK). The PC originated from pancreatic tissue of 12 months old Albino Swiss (Glasgow) rats. The pancreatic tissue was minced prior to culturing in serum-free medium and the PC emerged as a confluent monolayer after approximately 5 weeks in culture.

#### Human Dermal Fibroblasts (HDF)

HDF were cultured in Dulbecco’s modified Eagle’s medium (DMEM) with GlutaMAX™-I (Thermo Fisher Scientific, UK), supplemented with 10% heat inactivated FBS (Thermo Fisher Scientific, UK), 1% penicillin-streptomycin (Thermo Fisher Scientific, UK) and 1% Amphotericin B (fungizone®) (Thermo Fisher Scientific, UK).

#### Mesenchymal Stem Cells (MSC)

Human MSC were cultured in MSCBM™ Mesenchymal Stem Cell Basal Medium (MSCBM hMSC Basal Medium) (Lonza, UK) with necessary supplements (MSCGM hMSC SingleQuot Kit) (Lonza, UK). According to the manufacturer, the serum that was used neither promotes spontaneous differentiation of cells nor inhibits cell differentiation to osteogenic, chondrogenic or adipogenic lineages, when properly stimulated.

#### Vascular Smooth Muscle Cells (VSMC)

Collection, storage, and use of tissue and human aortic samples were performed in agreement with the Dutch Code for Proper Secondary Use of Human Tissue. VSMC were cultured in Medium 199 (Thermo Fisher Scientific, UK), supplemented with 10% heat inactivated FBS (Thermo Fisher Scientific, UK), 1% penicillin-streptomycin (Thermo Fisher Scientific, UK) and 1% Amphotericin B (fungizone®) (Thermo Fisher Scientific, UK) and 1% GlutaMAX™ (Thermo Fisher Scientific, UK). The VSMC originated from patients who had suffered a non-genetic abdominal aortic aneurysm. Specifically, the VSMC were isolated by a surgeon from healthy and aneurysm aorta of the same patient.

### 4.2. Extracellular vesicle (EV) isolation

Rat PC were cultured for the purpose of isolating MV and Exo. The PC were initially plated on Corning® T75 flasks (Merck, UK). Once the cells reached 70% confluency, they were transferred to bigger Corning® T150 flasks (Merck, UK). When the PC reached 70% confluency once again, they were used to prepare a 50 mL cell suspension in MV-free CMRL 1066 complete media. For this type of cell media, we used heat-inactivated FBS that was centrifuged at 16,000 ×g for 3 hours in order to remove potentially present MV within it. The cell suspension was then transferred to a Corning® HYPERFlask® cell culture vessel (Merck, UK), which was then completely filled with MV-free CMRL 1066 complete media. When the PCs reached 70% confluency, the culture media was collected and immediately transferred to Corning® 50 mL centrifuge tubes (Merck, UK) and centrifuged at 1000 ×g for 10 minutes to remove cell debris. The PC culture media supernatant was then transferred to a Corning® Easy-Grip round plastic storage bottle (Merck, UK) and stored at 4 °C, before it was used for MV and Exo isolation.

The PC culture media were split in Corning® 50 mL centrifuge tubes (Merck, UK) and centrifuged at 1000 ×g for 10 minutes to remove further cell debris. The supernatant was then carefully transferred to sterile Beckman Coulter® tubes (Beckman Coulter Life Sciences, USA) and centrifuged at 16000 ×g for 3 hours at 4 °C in an Avanti J-25 centrifuge (Beckman Coulter Life Sciences, USA) with a JA-25.50 fixed-angle aluminum rotor (Beckman Coulter Life Sciences, USA). The supernatant was used for Exo isolation (see next paragraph), while the MV pellets were resuspended in 1 mL sterile PBS. All of the resuspended MV were then transferred to a single Beckman Coulter® tube, in order to concentrate the MV sample. The single tube was then filled completely with sterile PBS and was centrifuged again at 16000 ×g for 3 hours. The supernatant was discarded and the MV pellet was resuspended in 1 mL sterile PBS. MV isolates were finally filtered twice with the use of MicroKros® Hollow Fiber Filters (Spectrum Laboratories, Inc., USA) 0.1 μm filters, in order to ensure removal of Exo contaminants. The MV isolates were finally stored at -20 °C.

The supernatant that contained the Exo was transferred to sterile Beckman Coulter® tubes and centrifuged at 120000 ×g for 3 hours at 4 °C. The supernatant was removed and the Exo pellets were resuspended in 1 mL sterile PBS. All of the resuspended Exo were then transferred to a single Beckman Coulter® tube, in order to concentrate the Exo sample. The single tube was then filled completely with sterile PBS and was centrifuged again at 120000 ×g for 3 hours. The supernatant was discarded and the Exo pellet was resuspended in 1 mL sterile PBS. Exo isolates were then filtered twice through 0.1 μm Minisart® Syringe Filters (Sartorius AG, Germany), in order to remove MV contaminants, which are larger than 0.1 μm in size. The Exo isolates were finally stored at -20 °C.

### 4.3. Flow cytometry

A drop of a series of Flow Cytometry Sub-Micron Size Reference Kit (Life Technologies, UK) size beads (0.2µm, 0.5µm, 1µm and 2µm) were added to 1 ml of PBS in 5mL round bottom polystyrene test tubes (BD Biosciences, UK) for flow cytometry and were run on a BD FACSVerse™ (BD Biosciences, UK) flow cytometer. 200 μL of the EV isolates were added directly in flow cytometry tubes and were run on the flow cytometer. The BD FACSuite™ (BD Biosciences, UK) software was used for the purposes of system setup, data acquisition, initial analysis and shutdown. The data that were collected were analysed with the use of FlowJo® Version 10.1 (FlowJo LCC, USA).

### 4.4. Western blot

The EV isolates were lysed in lysis buffer (50 mM Tris-HCl pH 7.4, 1% v/v Triton X100, 0.25% w/v sodium deoxycholate, 150 mM NaCl and 1 mM EDTA in distilled water). Protease inhibitors cOmplete™, Mini, EDTA-free (Merck, UK) and 1 mM phenylmethylsulfonyl fluoride (PMSF) (Merck, UK) were added at the time of lysis. The protein concentration of the EV was determined using the Quick Start™ Bradford protein assay according to manufacturer’s guidelines (BioRad, UK). In short, working reagent A was prepared by addition of 20 μL of reagent S to 1 mL of reagent A. A series of protein standards were then prepared (0.25 mg mL^-1^, 0.5 mg mL^-1^, 0.75 mg mL^-1^, 1 mg mL^-1^, 1.5 mg mL^-1^) from 10 mg mL^-1^ bovine serum albumin (BSA) in 0.15 M NaCl. Then, 5 μL of the prepared protein standards and lysed MV samples were loaded on a Corning® 96-well plate (Corning, UK). 25 μL working reagent A and 200 μL reagent B were also added. After 15 minutes at room temperature, the absorbance was read at 750 nm with the use of a GloMax®-Multi Detection System (Promega, UK), in order to determine the EV samples protein concentration. Then, 1µg of EV protein (15 μL) per lane was prepared in 5 μL NuPAGE™ LDS Sample Buffer (ThermoFisher Scientific, UK) and 2 μL NuPAGE™ Sample Reducing Agent (ThermoFisher Scientific, UK). The samples were heated at 70 °C for 10 minutes before they were loaded on a NuPAGE™ 12% gradient Bis-Tris Protein Gel (ThermoFisher Scientific, UK) and allowed to run at a constant current of 125 V, in the presence of 1x NuPAGE™ MOPS SDS Running buffer (Life Technologies, Inc., UK). The MV protein fractions were detected on the protein gel using Simple-Blue™ Safe Stain (ThermoFisher Scientific, UK). The sandwich method was then used to transfer the proteins from the gel to a methanol-activated polyvinylidene fluoride (PVDF) membrane (ThermoFisher Scientific, UK) at a constant current of 30 V for 1 hour and 30 minutes, in the presence of 1x NuPAGE™ transfer buffer (ThermoFisher Scientific, UK) with 20% methanol.

Immunoblotting was performed using standard protocols. Briefly, the PVDF membrane was washed 3 times with 1x StartingBlock™ Tris-buffered saline (TBS) (ThermoFisher Scientific, UK)–0.1% Tween 20 (TBST) and blocking was achieved with a 5% milk solution in TBST that was added to the membrane for 1 hour, while on a moving platform. Then the primary rabbit antibodies CD40 (1:500; cat. no. ab13545, Abcam, UK), Integrin β1 (1:500; cat. no. sc-8978, Santa Cruz Biotechnology, Inc., USA), Rab5b (1:300; cat. no. sc-598, Santa Cruz Biotechnology, Inc., USA), CD63 (1:1000; cat. no. EXOAB-CD63A-1, SBI, USA) and CD9 (1:500 dilution; cat. no.

ab92726, Abcam, UK) were added. The primary antibodies were allowed to bind overnight in at 4 °C, on a moving platform. The membrane was then washed 3 times with TBST and the secondary goat anti-rabbit IgG horseradish peroxidase (HRP)-conjugated antibody (1:5000; cat. no. 7074, Cell Signaling Technology, The Netherlands) was added and allowed to bind at room temperature for 1 hour, on a moving platform. Finally, the membrane was washed another 3 times with TBST, before equal volumes of substrates A and B of Enhanced Chemiluminescence (ECL) System (BioRad, UK) were added on top of the membrane to cover it and allowed to act on the antibodies for 30 seconds. The membrane was then developed with the use of a G:BOX (Syngene, UK).

### 4.5. Senescence-associated β-galactosidase (SA β-gal) assay

The cells were stained for SA β-gal 72 hours after plating 50000 cells per well, in 6-well plates. Initially, the Senescence Cells Histochemical Staining Kit (Merck, UK) was used according to the manufacturer’s guidelines. Briefly, the cell media were aspirated and the cells were washed twice with PBS. The fixation buffer was then added and the cells were incubated for 7 minutes at room temperature. The cells were washed 3 times with PBS and 1.5 mL of the staining mixture (for 10 mL mixture: 8.5 mL milli-Q water, 1 mL staining solution, 125 μL Reagent B, 125 μL Reagent C and 250 μL X-gal Solution, filtered through 0.2µm pore size membrane) was added. The cells were incubated in the staining mixture overnight at 37 °C without CO2, pH 6, until they were stained blue. The cells were then washed three times with PBS and then they were stained with 1.43µM DAPI (Thermo Fisher Scientific, UK), diluted in PBS, at room temperature for 1 hour on a moving platform and covered from light. The DAPI staining step was followed by three washes with PBS. The cells were finally covered with 2 mL 70% glycerol solution diluted in milli-Q water. Visualisation was achieved with brightfield microscopy, and pictures were taken using a Zeiss Axio Observer microscope (Carl Zeiss Ltd., UK). Total cell number and cells positive for SA β-gal were scored. At least 1000 cells were counted per well.

### 4.6. γH2A.X staining

The cells were stained for γH2A.X and DAPI 72 hours after plating. The cell media were initially aspirated and the cells were washed 2 times with PBS. The cells were then fixed with 100% methanol (Merck, UK) at room temperature for 5 minutes. The cells were washed another 2 times and permeabilised with 0.1% Triton™ X-100 (Merck, UK) diluted in PBS and incubated at room temperature for 5 minutes. The cells were washed another 2 times with PBS before addition of blocking solution. The blocking solution consisted of 1% Bovine Serum Albumin, 10% goat serum, 0.3 M glycine in 0.1% PBS-Tween® 20 (All Merck, UK). Incubation took place at room temperature for 5 minutes. The primary rabbit antibody anti-γH2A.X (phospho S139) (cat. No. ab2893, Abcam, UK) was added in a concentration of 1µg mL^-1^ and the cells were incubated at 4°C on a moving platform overnight. After removal of the primary antibody and 3 washes with PBS, the secondary goat anti-rabbit Alexa Fluor® 488 antibody (1:1000; cat. no. ab150081, Abcam, UK) was added and allowed to bind at room temperature for 1 hour, on a moving platform. The secondary antibody was then removed and the cells washed thrice with PBS. Finally, the cells were stained with 1.43µM DAPI (Thermo Fisher Scientific, UK), diluted in PBS, at room temperature for 1 hour on a moving platform, covered from light. The DAPI staining step was followed by three washes with PBS and the cells were covered with 70% glycerol diluted in milli-Q water. Pictures were then taken with fluorescence microscopy, using a Zeiss Axio Observer microscope (Carl Zeiss Ltd., UK). Total cell number and the number of cells expressing cytoplasmic chromatin fragment (CCF) events were recorded. At least 1000 cells were counted per well.

### 4.7. CDKN2A and CDKN1A expression

#### RNA extraction

Cell samples (HDF, MSC or VSMC) were extracted before stress (wounding or uraemic serum addition) and 72 hours after stress in 1 mL TRIzol™ reagent (Thermo Fisher Scientific, UK) with the use of a cell scraper. The samples were stored at -20 °C overnight. The samples were then centrifuged at 12000 ×g for 10 minutes at 4 °C and the supernatants were transferred to fresh tubes on ice. Per 1 mL of TRIzol™ used, 200 μL of chloroform (Thermo Fisher Scientific, UK) were added. Hence, 200 μL chloroform were used. The samples were mixed by hand for 15 seconds and incubated at room temperature for 3 minutes. They were then centrifuged at 12500 ×g for 15 minutes at 4 °C. The aqueous phase was transferred to fresh tubes on ice. For one volume of aqueous phase, an equal volume of 95% ethanol (Merck, UK) diluted in milli-Q water was added.

The RNA Clean & Concentrator™-25 kit (Zymo Research, USA) was then used. Briefly, the samples were transferred to Zymo-Spin™ IIC columns in collection tubes and spin at 14000 ×g for 30 seconds. After the flow-through was discarded, the column was pre-washed with 400 μL RNA wash buffer, centrifuged at 14000×g for 30 seconds and the flow-through was discarded. DNase I mix (75 μL DNA digestion buffer and 5 μL DNase I per sample) was added to each column and the columns were then incubated at room temperature for 15 minutes.

400 μL RNA Prep buffer were added and the columns were centrifuged at 14000 ×g for 30 seconds. The flow-through was discarded. Then, 700 μL RNA wash buffer were added and the columns were centrifuged at 14000 ×g for 30 seconds. The flow-through was discarded. This step was repeated once more with 400 μL RNA wash buffer and centrifuged for 2 minutes this time. The flow-through was discarded and each sample was transferred to a new Eppendorf tube. 15 μL DNase/RNase-free water were added to each column and centrifuged at 14000 ×g for 30 seconds. This step was repeated once more and the 30 μL total flow-through was collected and placed on ice for 15 minutes.

Finally, 1.6 μL of the RNA extracts were used for RNA measurement with the use of the NanoDrop™ 2000 Spectrophotometer (Thermo Fisher Scientific, UK). The RNA samples were then stored at -80 °C.

#### RT-PCR

2.5 μL random primers (3µg μL^-1^) (Thermo Fisher Scientific, UK) and 2.5 μL Deoxynucleotide mix (10 mM) (Thermo Fisher Scientific, UK) were added to 125 ng RNA per sample. Nuclease-free water (Thermo Fisher Scientific, UK) was added to a final volume of 19 μL. The samples were mixed well by pipetting and incubated at 67 °C for 5 minutes. The tubes were then placed on ice before adding 16 μL of the prepared pre-mix, including per well 8 μL 5X first-strand buffer, 2μL RNaseOUT™ recombinant ribonuclease inhibitor (40 U μL^-1^), 4 μL 0.1M DTT and 2 μL SuperScript® II Reverse Transcriptase (200 U μL^-1^). The samples were then mixed well by pipetting.

#### qPCR

The qPCR mastermix was prepared following the manufacturer’s guidelines (Thermo Fisher Scientific, UK). Briefly, each reaction contained 5 μL TaqMan™ Universal PCR mastermix II (no UNG), 3.5 μL nuclease-free water, 0.5 μL 20X TaqMan™ Assays qPCR primers (all Thermo Fisher Scientific, UK) (Table 2) and 1 μL of product of RT-PCR reaction. Relative quantity of RNA was analysed using comparative threshold (Ct) method. Negative controls included a no template control (no RNA at cDNA generation step) and an amplification control (no cDNA added).

### 4.8. 5-bromo-2’-deoxyuridine (BrdU) labelling

Stress assays (wounding or uraemic serum addition) were performed in Corning® 6-well plates (Merck, UK). The HDF, MSC and VSMC (healthy and aneurysm) were counted and used immediately for plating on 6-well plates as monocultures. After allowing for overnight attachment in the incubator at 37 °C and 5% CO2, a scratch wound was performed with a cell scraper for the wound healing assays. The media were then gently removed and replaced with fresh MV-free DMEM media. For the uraemic serum assays, the media were gently removed and replaced with fresh MV-free DMEM media, 10% uraemic serum and MV. MV were administered at a dose of 0.1 ng μL^-1^ RNA per well. For control, the wells were replaced with MV-free media containing equal volume of PBS, instead of MV.

The cells were stained with BrdU before stress and 72 hours after stress. Initially, the 5-Bromo-2’-deoxy-uridine Labelling and Detection Kit II (Roche, UK) was used per manufacturer’s protocol. The cell media were aspirated and the cells were washed twice with PBS before the BrdU labelling medium (1:1000 dilution in MV-free complete media, filtered through 0.2µm pore size membrane) was added for the labelling of DNA. The cells were incubated with the BrdU labelling medium at 37 °C for 30 minutes. They were then washed 3 times with washing buffer (diluted 1:10 in milli-Q water). The cells were then fixed in ethanol fixative (30 mL 50 mM glycin solution and 70 ml absolute ethanol, pH 2.0) for 10 minutes at -20 °C. The cells were then washed another 3 times with washing buffer and the Anti-BrdU solution was added and allowed to act at 37 °C for 30 minutes. The cells were washed 3 times again and the Anti-mouse-Ig-AP solution was added to the cells for 30 minutes at 37 °C. Finally, the cells were washed 3 times with washing buffer and freshly prepared colour substrate buffer solution (100 mM Tris HCl, 100 mM NaCl, 50 mM MgCl2, pH 9.5) was added at room temperature for 15 minutes before it was washed with washing buffer 2 times as well. The cells were then covered with 70% glycerol diluted in milli-Q water.

The cells were visualised with brightfield microscopy, using a Zeiss Axio Observer microscope (Carl Zeiss Ltd., UK). Total cell number and cells positive for BrdU were counted. At least 1000 cells were counted per well.

### 4.9. Real time cell analysis (RTCA)

Wound healing and uraemic serum assays were performed in E-Plate VIEW 96 (ACEA Biosciences, Inc., USA), which is a specific type of 96-well plate, fused with gold microelectrodes that are capable of detecting the presence of adherent cells. Cell growth was recorded with the xCELLigence® RTCA MP (ACEA Biosciences, Inc., USA). 100 μL DMEM media per well were added and the plate was placed in the RTCA instrument in the incubator (37 °C, 5% CO2), to measure the background impedance. Additionally, a cell suspension in complete DMEM was prepared for HDF and MSC (or VSMC). HDF and MSC (or VSMC) were counted and 5000 cells were plated as a co-culture, at a ratio of 5:1 HDF to MSC (or HDF to VSMC). The plate was filled to 200 μL per well with addition of the mixed cell suspension and the cells were allowed to sediment for 30 minutes in the tissue culture hood. The plate was then placed in the RTCA instrument in the incubator for overnight incubation. 24 hours later, after allowing for overnight attachment, a scratch wound was performed with a 20 μL sterile pipette tip for each well for the wound healing assays. For the uraemic serum assays, the media were gently removed and replaced with 150µL fresh MV-free DMEM media, 20µl of uraemic serum (10% final concentration) and 30µL pathfinder cell MVs (or Exo) (0.1 ngµL^-1^ of total RNA). The media were then gently removed and replaced with 170 μL fresh MV-free DMEM media and 30 μL PC MV (or Exo) (0.1 ng μL^-1^ of total RNA). For control, the wells were replaced with media containing equal volume of PBS instead of MV. The plate was returned to the RTCA instrument in the incubator and wound closure was recorded during a three-day period. Wells containing no cells and only media or media with MV were also used as controls.

Cell proliferation assays in the absence of strerss were also performed in E-Plate VIEW 96 and cell growth was recorded with the xCELLigence® RTCA MP. 100 μL MV-free DMEM media per well were added and the plate was placed in the RTCA instrument in the incubator (37 °C, 5% CO2), to measure the background impedance. Additionally, a cell suspension in MV-free DMEM media was prepared for HDF and MSC. HDF and MSC were counted and 5000 cells were plated as a co-culture, at a ratio of 5:1 HDF to MSC. The plate was filled to 200 μL per well with addition of the mixed cell suspension and MV (0.1 ng μL^-1^) or a similar volume of PBS instead of MV as a control. The cells were allowed to sediment for 30 minutes in the tissue culture hood. The plate was then placed in the RTCA instrument in the incubator and cell proliferation was monitored for 72 hours. Wells containing no cells and only media or media with MV were also used as controls.

Cell migration assays in the absence of stress were performed in CIM-Plate 16 (ACEA Biosciences, Inc., USA), which are a specific type of 16-well plate comprising of an upper chamber with a microporous membrane (8 μm pore diameter) and a bottom chamber. Gold electrodes on the underside of the upper membrane detect the presence of cells migrating through the pores. Cell migration was recorded with the xCELLigence® RTCA DP (ACEA Biosciences, Inc., USA). The bottom chamber of the 16-well plate was completely filled with MV-free DMEM. The upper chamber was locked on top of the bottom chamber and it was filled with 50 μL MV-free DMEM media. The fully assembled plate was placed in the RTCA instrument in the incubator (37 °C, 5% CO2), to measure the background impedance. A cell suspension in MV-free DMEM was prepared for HDF and MSC. Cells were then counted and the two cell types were mixed at a 5:1 ratio of HDF to MSC. 5000 cells were then added to the upper chamber at a total volume of 170 μL per well, along with 30 μL of MV (0.1 ng μL^-1^) or a similar volume of PBS instead of MV as a control. Cells were allowed to sediment for 30 minutes in the incubator. The plate was then placed in the RTCA instrument in the incubator and cell migration was monitored for 72 hours. This experiment was also repeated with addition of a similar dose of MV (or PBS as a control) at the bottom chamber. Wells containing no cells and only media or media with MV were also used as controls.

The RTCA normalized cell index over time data, which were collected, were analysed with the use of a slope analysis by the RTCA software version 1.2.1 (ACEA Biosciences, Inc., USA).

### 4.10. Time lapse fluorescence microscopy

Wound healing assays were performed in Corning® 6-well plates (Merck, UK). HDF and MSC were individually labelled with cell tracking dyes eBioscience™ CFSE (Thermo Fisher Scientific, UK) and PKH26 (Merck, UK) respectively. Optimised protocol was used for both cell types. After preparing serum-free cell suspension, the fluorescent cell tracking dyes were added at a final concentration of 5 μM for both dyes and the cells were incubated in the dark at 37 °C for 10 minutes. Labelling was stopped by addition of 5 volumes of cold complete media and incubation on ice for a further 5 minutes. The cells were then washed 3 times with complete media to remove excess of the cell tracking dyes.

The cells were counted and used immediately for plating on 6-well plates as a co-culture of 5:1 ratio HDF to MSC. After allowing for overnight attachment in the incubator at 37 °C and 5% CO2, a scratch wound was performed with a cell scraper. The media were then gently removed and replaced with fresh MV-free DMEM media. MV were administered at a dose of 0.1 ng μL^-1^ RNA per well. For control, the wells were replaced with MV-free media containing equal volume of PBS, instead of MV.

Pictures were taken every 12 hours over a 3-day period, with fluorescence microscopy, using a Zeiss Axio Observer microscope (Carl Zeiss Ltd., UK). The number of cells migrating into the site of wounding during repair were counted.

## ACKNOWLEDGEMENTS

N.P. was supported by a studentship award from the Biotechnology and Biological Sciences Research Council (BBSRC). N.P. and D.M. were supported by endowment awards from NHSGGC. A.J., B.M. and L.S. were supported by the public-private partnership RegMed-XB. We thank Sarah Buchanan for technical assistance with experiments.

## AUTHOR CONTRIBUTIONS

P.G.S., N.P., C.S. and L.S. conceptualised the study; N.P. conducted experiments and analysed data; D.M. contributed to study design. N.P. and D.M. EV isolation and characterisation; A.M.G.J. and B.M. VSMC and AAA VSMC isolation and preparation. N.P., P.G.S. and C.S. wrote the manuscript; All authors read and edited the manuscript. Project administration, P.G.S. and C.S.; Funding acquisition, P.G.S., C. S., D.M. and N.P.

## DATA AVAILABILITY STATEMENT

All the data that support the figures and the other findings are available upon request to the corresponding author.

