## Supplemental for "Microvesicle-Mediated Tissue Regeneration Mitigates the Effects of Cellular Ageing"

### Supplementary Figures:

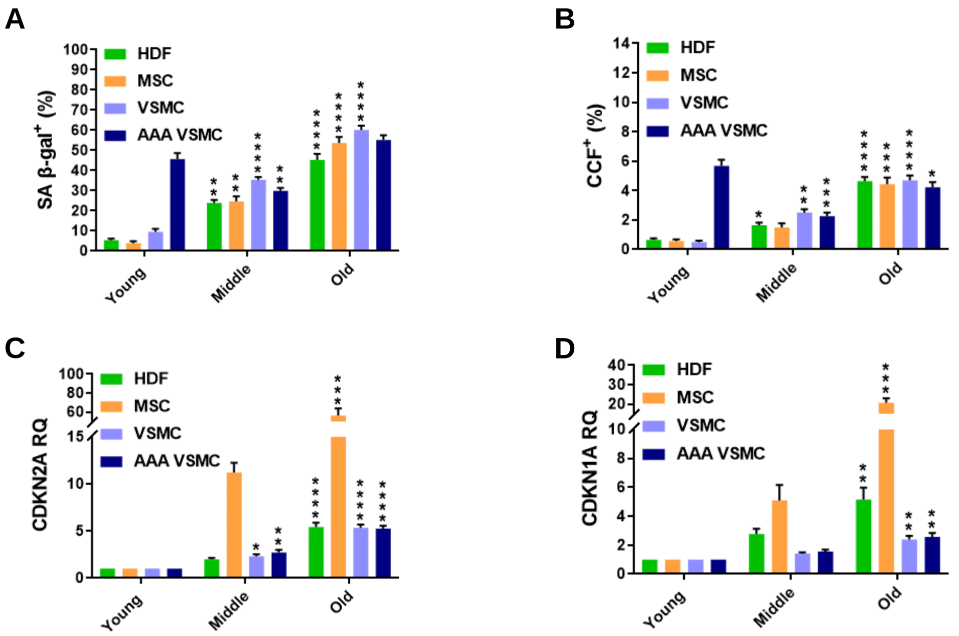


**Figure S1: Ageing markers track cellular ageing *in vitro*.**

**(A)** Percentage of SA β-gal$^{+}$ cells with increasing cellular age. Data are presented as mean ± SD; One-way ANOVA with Dunnett's test (N=3).

**(B)** Percentage of CCF$^{+}$ cells with increasing cellular age. Data are presented as mean ± SD; One-way ANOVA with Dunnett's test (N=3).

**(C)** CDKN2A expression with increasing cellular age. Data are presented as mean ± SD; One-way ANOVA with Dunnett's test (N=3).

**(D)** CDKN1A expression with increasing cellular age. Data are presented as mean ± SD; One-way ANOVA with Dunnett's test (N=3).

*p≤0.05, **p≤0.01, ***p≤0.001, ****p≤0.0001.

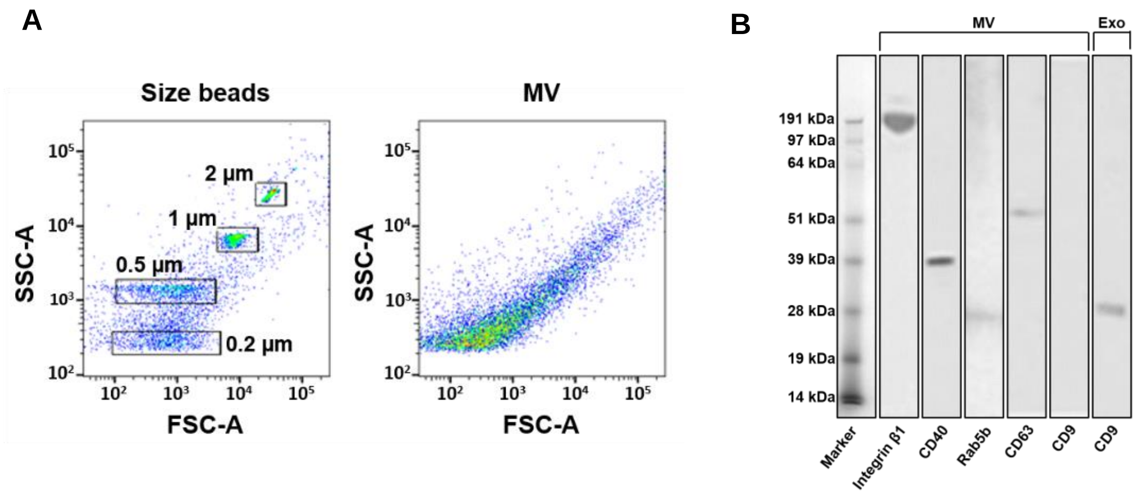


**Figure S2: Characterisation of PC-derived MV and Exo isolates.**

**(A)** Size profile of PC MV characterised by forward scatter (FSC-A) vs. side scatter (SSC-A) flow cytometry in comparison to a series of beads of known size, including 0.2 µm, 0.5 µm, 1 µm and 2 µm. **(B)** Surface markers of PC MV and Exo isolates. The representative immunoblots above show the presence of MV markers (Integrin β1, 138 kDa; CD40, 43 kDa; Rab5b, 25 kDa and CD63, 53 kDa) and the absence of exosome marker (CD9, 28 kDa) in the MV isolates. Exo isolates express only CD9.

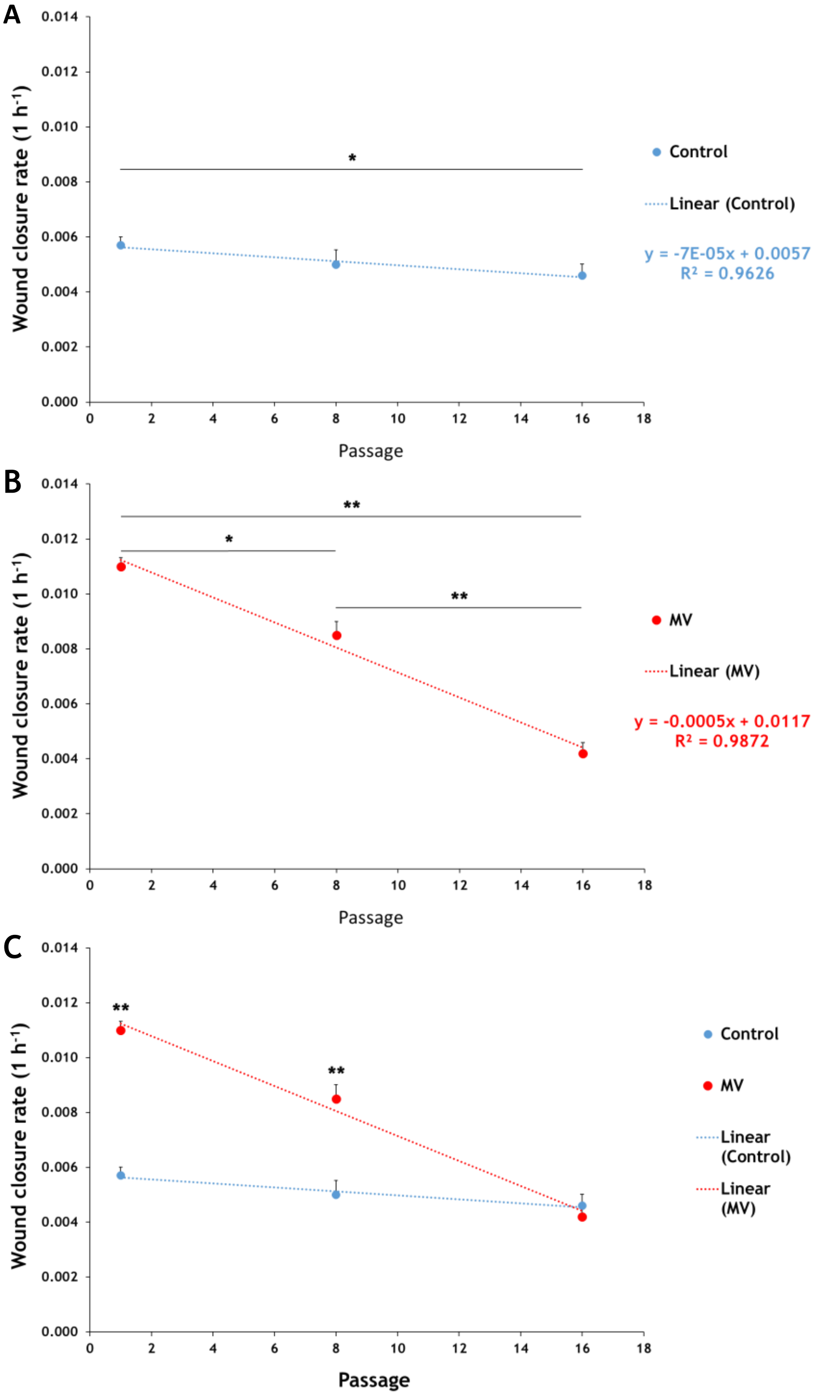


**Figure S3: Wound closure rate with increasing cellular age.**
**(A)** Physiological wound closure rate in HDF-MSC mixed cultures of increasing cellular age. Data are presented as mean ± SD; one-way ANOVA with Tukey's HSD test (N=3).
**(B)** Effect of MV administration during wound repair in HDF-MSC co-cultures of increasing cellular age. Data are presented as mean ± SD; one-way ANOVA with Tukey's HSD test (N=3).
**(C)** Combined. MV administration (red). PBS administration instead of MV (blue). Data are presented as mean ± SD; t test (N=3).
*p<0.05, **p<0.01.

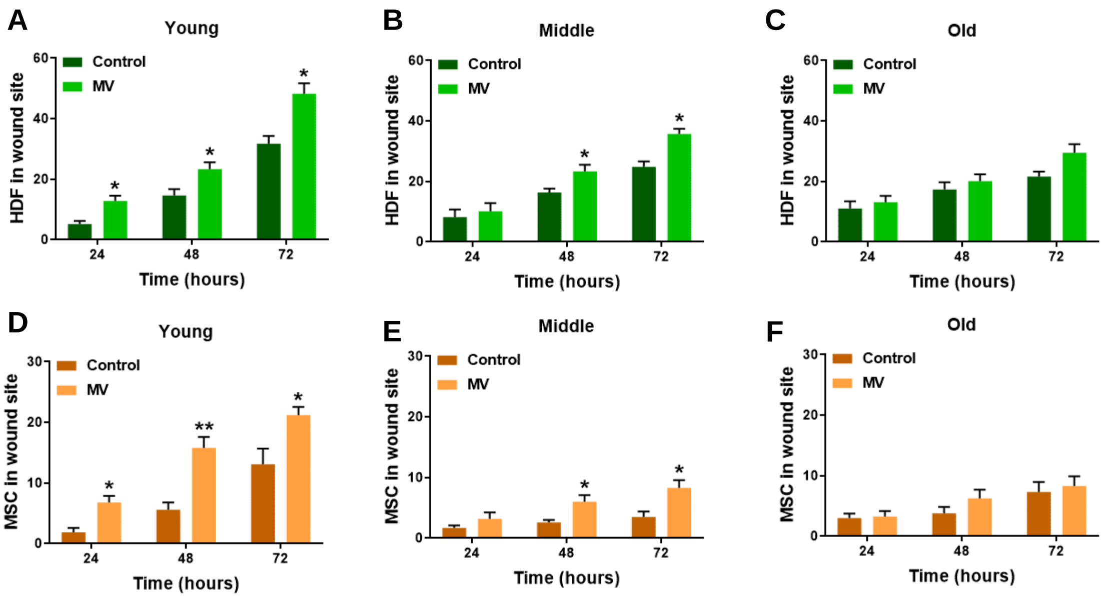


**Figure S4: Quantification of HDF and MSC migration into the wound site.**
Effect of MV administration on HDF migration into the wound site; 24, 48 and 72 hours after wounding in **(A)** young, **(B)** middle, and **(C)** old HDF-MSC co-cultures. Data are presented as mean ± SD; t test (N=3).
Effect of MV administration on MSC migration into the wound site; 24, 48 and 72 hours after wounding in **(D)** young, **(E)** middle, and **(F)** old HDF-MSC co-cultures. Data are presented as mean ± SD; t test (N=3).
*p≤0.05, **p≤0.01.

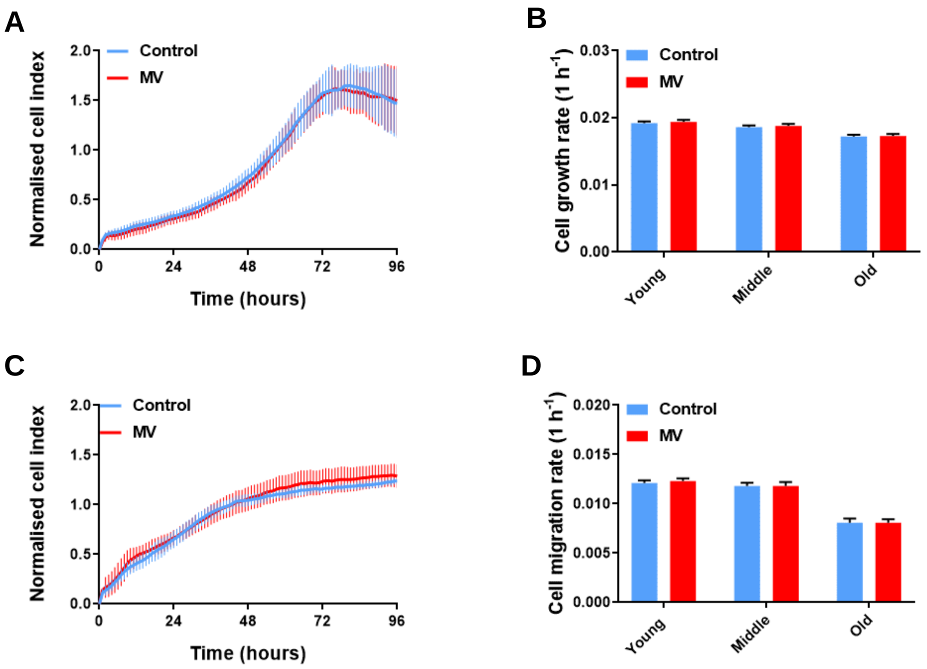


**Figure S5: The proliferative and migratory capacity of HDF-MSC co-cultures declines with cellular age, while MV do not affect these processes in the absence of cellular damage.**

**(A)** RTCA proliferation assay on a young HDF-MSC co-culture. Data are presented as mean ± SD (n=6).

**(B)** Cell growth rate of HDF-MSC co-cultures with cellular age. Data are presented as mean ± SD; t test (N=3).

**(C)** RTCA migration assay on a young HDF-MSC co-culture. Data are presented as mean ± SD (n=6).

**(D)** Cell migration rate of HDF-MSC co-cultures with cellular age. Data are presented as mean ± SD; t test (N=3).

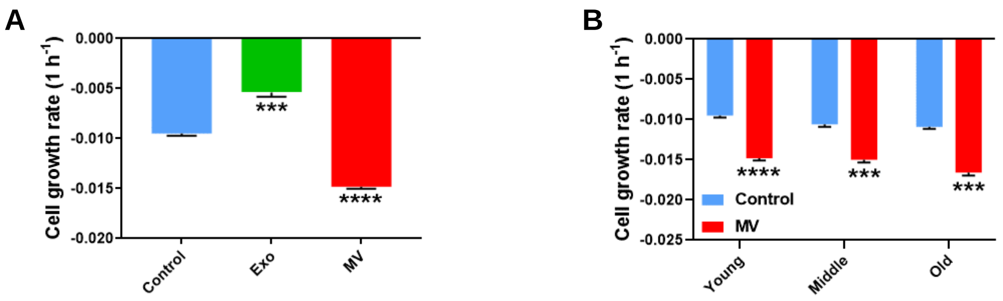


**Figure S6: Rate of cell growth in HDF-MSC genotoxicity assays from 0 to 24 hours.**
**(A)** Cell growth rate of young HDF-MSC co-cultures treated with MV and Exo, 24 hours after uraemic serum addition. Data are presented as mean ± SD; One-way ANOVA with Dunnett's test (N=3).
**(B)** Effect of MV administration during the first 24 hours of genotoxic stress in HDF-MSC co-cultures of increasing cellular age. Data are presented as mean ± SD; t test (N=3).
***p≤0.001, ****p≤0.0001.

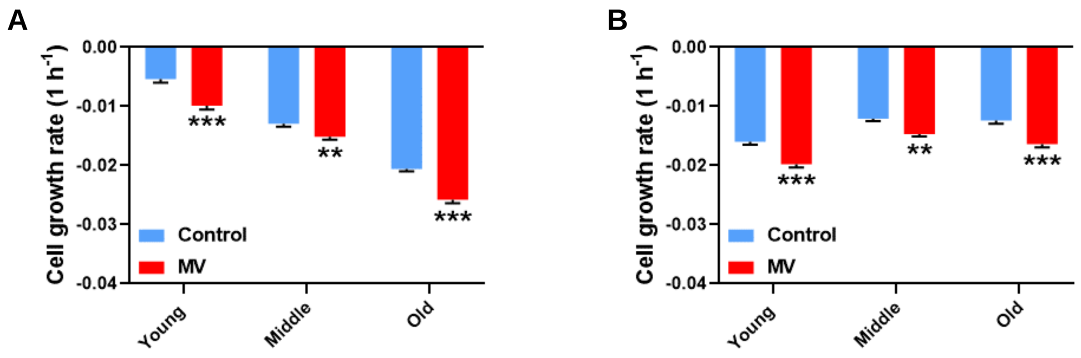


**Figure S7: Rate of cell growth in HDF-VSMC and HDF-AAA VSMC genotoxicity assays from 0 to 24 hours.**

**(A)** Effect of MV administration during the first 24 hours of genotoxic stress in HDF-VSMC co-cultures of increasing cellular age. Data are presented as mean ± SD; t test (n=3).

**(B)** Effect of MV administration during the first 24 hours of genotoxic stress in HDF-AAA VSMC co-cultures of increasing cellular age. Data are presented as mean ± SD; t test (n=3).

**p≤0.01, ***P≤0.001.
